# Rapid temporal adaptation structures tolerance to toxic cyanobacteria in a natural population of the water flea *Daphnia*

**DOI:** 10.1101/2025.02.28.640829

**Authors:** Maxime Fajgenblat, Emma Gouwy, Manon Coone, Rafaela Almeida, Alice Boudry, Kiani Cuypers, Edwin van den Berg, Isabel Vanoverberghe, Luc De Meester, Ellen Decaestecker

**Author notes:** Corresponding author: Maxime Fajgenblat. Shared first authorship (Maxime Fajgenblat, Emma Gouwy). Shared senior authorship (Luc De Meester, Ellen Decaestecker).

## Abstract

Cyanobacteria blooms pose a substantial threat to freshwater systems globally. While zooplankton grazers such as *Daphnia* can have an important role in suppressing cyanobacteria blooms, cyanobacteria can adversely impact *Daphnia* fitness and even kill them. Earlier work has shown an evolutionary increase in tolerance to cyanobacteria across years and strong genotype x genotype interactions determining the interaction between *Daphnia* and the cyanobacterium *Microcystis*. Here, we test the hypothesis that *D. magna* can adapt during one growing season to changes in dominant strains of *Microcystis*. Over two consecutive years, we collected *D. magna* clonal lineages and *Microcystis* strains from a single pond early and late in the growing season and we assessed whether *Daphnia* survival differed when exposed to *Microcystis* strains from either the same or a different time point within the growth season. Our findings reveal important *Daphnia* genotype x *Microcystis* genotype interactions, with *Daphnia* survival being higher when exposed to *Microcystis* from the same time point than when exposed to *Microcystis* of a different time point. Our results extend earlier findings to variation within one single natural system and growth season, and suggest an important impact of rapid (co)evolutionary dynamics shaping the tolerance of zooplankton grazers to cyanobacteria.

## Introduction

Cyanobacteria blooms are increasingly common across the globe because of rising temperatures and nutrient levels (Kosten et al., 2012; Meerhoff et al., 2022). Cyanobacteria blooms can exert substantial pressure on ecosystem functioning due to their effect on the fitness of many organisms (Pimentel et al., 2001). They reduce nutrient and light availability for submerged macrophytes, can typically not be grazed because they form colonies, are generally a low-quality food source for grazers, and can produce toxins (Carton, 2005; Gray et al., 2014; Paerl et al., 2011; Sommer et al., 2012). As such, they can shift shallow lakes to a turbid state dominated by cyanobacteria, that is difficult to control by grazers (Hansson et al., 2007; Scheffer & Van Nes, 2007). In temperate regions, cyanobacteria blooms typically occur in summer (PEG model; Sommer & Maciej Gliwicz, 1986). Higher temperatures, along with nutrient release from the sediment (Paerl & Huisman, 2008) and decreased zooplankton grazing caused by fish predation (Lampert & Sommer, 2007) lead to an increased fraction of cyanobacteria in the phytoplankton community.

The water flea *Daphnia* plays a pivotal role in many standing freshwaters, as they are key grazers on phytoplankton and the preferred prey of fish (Miner et al., 2012). *Daphnia* can often exert a strong top-down control on phytoplankton (Gianuca et al., 2016; Jeppesen et al., 1996), and can even make the difference between a cyanobacteria bloom and a clear-water state (Sarnelle & Wilson, 2005; Scheffer et al., 2001). Yet, many studies have reported that *Microcystis* can kill *Daphnia* due to the production of toxins or feeding inhibition and starvation, for instance through its low nutritional quality, the production of digestive protease inhibitors and colony formation (Agrawal et al., 2001; Ghadouani et al., 2004; Lürling, 2003; Macke et al., 2017; Trubetskova & Haney, 2006; von Elert et al., 2003). Consequently, while *Daphnia* can serve as an efficient grazer of *Microcystis*, it can also be vulnerable to it. This leads to a large variation in results on top-down control of cyanobacteria blooms by *Daphnia*, as reported in field-based and experimental work (e.g. Benndorf & Henning, 1989; de Bernardi & Guissani, 1990; Dejenie et al., 2009; Gulati et al., 2008; Sarnelle & Wilson, 2005; Søndergaard et al., 2008). Several studies indicate that *Daphnia* displays inducible responses (which can be transferred through maternal effects; Gustafson et al. 2005) and can even genetically adapt when exposed to cyanobacteria (e.g. through selection on digestive proteases; Schwarzenberger et al., 2020), which may contribute to the variation in empirical outcomes. Lemaire et al. (2011) experimentally showed both variation in tolerance to *Microcystis* among *Daphnia* genotypes and variation in overall toxicity of *Microcystis* strains, but also striking genotype x genotype interactions that make the observed survival of *Daphnia* when exposed to *Microcystis* dependent on the specific combination of *Daphnia* and *Microcystis* genotypes. Such genotype x genotype interactions are often observed in host-parasite interactions and can lead to rapid local adaptation and Red Queen dynamics (Capaul & Ebert, 2003; Decaestecker et al., 2007; Lambrechts et al., 2006). A resurrection study has indeed shown that *Daphnia* in Lake Constance evolved higher tolerance to cyanobacteria as the lake eutrophied (Hairston et al., 1999, 2001) and subsequently lost this adaptation as nutrient concentrations in the lake became lower again (Isanta-Navarro et al., 2021). In line with these results, Sarnelle and Wilson (2005) observed that *D. pulicaria* populations from eutrophied ponds are significantly more tolerant to cyanobacteria than *D. pulicaria* populations from less eutrophic ponds.

Because of their role as key grazers of phytoplankton, the capacity of *Daphnia* populations to evolve enhanced tolerance to cyanobacteria can have important ecological consequences. By combining field monitoring with genotyping and laboratory experiments, Schaffner et al. (2019) showed that *D. mendotae* clones dominant in summer are better able to feed on low quality algae communities with a high proportion of cyanobacteria and that this evolutionary change increased their abundance and thus the potential for top-down control of phytoplankton. Hegg et al. (2022) did not find evidence for increased tolerance to cyanobacteria when comparing clones obtained before and after a bloom. In this experiment of Hegg et al. (2022), however, standard laboratory strains were used rather than local strains of *Microcystis*. If genotype x genotype interactions are important and lead to patterns of local (co)adaptation, the use of laboratory strains of *Microcystis* might fail to reveal the resulting eco-(co)evolutionary dynamics.

In the present study, we set out to test whether adaptation of *Daphnia* populations to changes in *Microcystis* strain composition within a single growing season can be an important aspect of eco-(co)evolutionary dynamics that determine the outcome of the interaction between *Daphnia* and *Microcystis*. More specifically, we tested whether a *D. magna* population inhabiting a shallow lake can genetically track changes in *Microcystis* strain composition through rapid evolution as a bloom develops in spring. Over two consecutive years, we harvested *Daphnia* and *Microcystis* from two time points in the growing season. We subsequently performed a reciprocal exposure experiment where *Daphnia* clonal lineages were exposed to *Microcystis* strains from either the same or the other time point. We tested the hypotheses that (i) *Daphnia* genotype x *Microcystis* genotype interactions occur with the capacity of *Daphnia* to cope with *Microcystis* within the set of genotypes and strains isolated from a single lake, and (ii) *Daphnia* clonal lineages are more tolerant to *Microcystis* strains isolated during the same moment in the growing season than to *Microcystis* strains isolated from a different moment in the growing season.

## Material and methods

### Isolation of Daphnia and Microcystis

We isolated twelve clonal lineages of *D. magna* and twelve isolates of *Microcystis* sp. from Langerodevijver, a manmade pond located in a nature reserve in Belgium (50°49’42.9”N; 4°38’23.8”E). To capture the interannual and seasonal dynamics within Langerodevijver, isolation took place in two consecutive years (2018 and 2019) during two distinct time periods that we will further refer to as ″early″ and ″late″ in the growing season: end of April 2018, end of May 2018, end of April 2019 and mid to end of June 2019 (Table S1). These time points were chosen because of the temporal dynamics of phytoplankton, cyanobacteria and different *Daphnia* species in this pond (see Fig. S1), where *Daphnia* appears in samples around mid-March and disappears from the pond around June, presumably because of predation by young-of-the-year fish. Cyanobacteria are still at low densities, and generally start to bloom after the disappearance of *D. magna*; as a result all isolations of *Daphnia* and *Microcystis* were carried out before the peak annual *Microcystis* sp. summer bloom. At each of the four sampling moments, three *D. magna* females were isolated to establish clonal lineages and a water sample was taken and stored at 4°C for isolation of *Microcystis* sp. strains.

### *Daphnia magna* cultures

*D. magna* individuals were individually placed in 150 mL glass jars filled with filtered tap water (Greenline e1902 filter) to establish clonal lineages, which were maintained under standardized laboratory conditions prior to the experiment (16:8 h light–dark cycle, constant ambient temperature of 19 ± 1°C, fed three times a week 200×10^3^ cells/mL of *Chlorella vulgaris*, and medium refreshed once a month). Before the start of the experiment, all clonal lines were kept under optimal conditions for at least two generations to exclude interference of maternal effects and to ensure that the observed responses were due to genetic effects. Animals were kept individually in 50 mL of aged tap water (16:8 h light–dark cycle, constant ambient temperature of 19 ± 1°C, fed three times a week 200×10^3^ cells/mL of *Chlorella vulgaris*, and medium refreshed once a week). Both the maternal generation and the generation of experimental animals were initiated by inoculating one neonate <24 h old from the second or third brood.

### *Microcystis* sp. cultures

For the isolation of *Microcystis* sp., three droplets of sampled water were placed on a glass microscope slide and placed under an SZX10 stereomicroscope (Olympus, Tokyo, Japan). Single colonies were then picked out using the following mouth-pipetting technique: the tip of a glass Pasteur pipette was heated using a Bunsen flame and then extended using forceps. This extended pipette was then connected to an autoclaved flexible, plastic tube. The other end of the tube was placed in the mouth of the person isolating the cells. Single colonies were identified under the microscope and carefully pipetted up by exerting gentle suction on the plastic tube end. The isolated colony was then washed three times in modified Wright’s Crytophyte (WC) medium (i.e. without Tris and a fivefold of NaNO_3_ and K_2_HPO_4_, supplemented with vitamins), known to promote *Microcystis* sp. growth (Guillard & Lorenzen, 1972). The single *Microcystis* sp. colony was then placed in 2 mL of this medium in a 24-well plate. Plates were covered with parafilm to avoid contamination and placed on a shaker under a 16:8 h light cycle at 21 ± 1°C. Additionally, plates were covered with a neutral density light filter to reduce light intensity and to stimulate *Microcystis* sp. growth. Further manipulation of the cyanobacterial cultures was performed under the flow hood. Once the 2 mL culture showed a healthy green colour, it was added to 5 mL of fresh medium in a 6-well plate under the flow hood. The empty well was rinsed thoroughly, and the new plate was covered with parafilm and placed on a shaker. Once the cultures were dense, three cultures per timepoint were selected and individually transferred to a 1 L glass culture bottle (Düran) under aeration, with the bottle caps fitted with 0.22 µm Millipore air filters. At the aeration system’s input and output to supply CO_2_ and to remove oxygen, a second 0.22 μm filter was used to prevent (cyano)bacterial contamination. To maintain regular mixing and prevent algal precipitation, magnetic stirrers were used. The algal culture cell density was assessed using flow cytometry on the FACS Verse cell sorter (Biosciences). Prior to cell counting, colonies were separated into individual cells by vigorously shaking the vials using a vortex mixer.

### Experimental set-up

The experiment was conducted in February 2021. *D. magna* clonal lineages were administered *Microcystis* sp. strains from either the same (further referred to as “contemporal”) or a different timepoint (further referred to as “allotemporal”) within the same year (Fig. S2). We did this for *D. magna* clonal lineages and *Microcystis* sp. strains from Langerodevijver isolated in two consecutive years, enabling us to assess the repeatability of the observed pattern over time. We considered 72 combinations of *Daphnia* clonal lineages and *Microcystis* strains (2 years x 2 *Daphnia* populations x 3 *Daphnia* clonal lineages per population exposed to 2 *Microcystis* populations x 3 *Microcystis* strains per population) as experimental units, each including six experimental animals. One *Daphnia* clonal lineage (*4b*) was excluded because it did not produce sufficient brood. Two other clonal lineages (*1c* and *4a*) did not produce sufficient brood to use six experimental animals, leading to 11 experimental units that only contained four or five individuals. Consequently, 66 out of the 72 experimental units with a total of 380 experimental *Daphnia* individuals were effectively executed. The *Daphnia* individuals were cultured for a total of 12 days, since this duration has been shown to highlight clonal differences in sensitivity to cyanobacteria (Macke et al., 2017). Starting from day one of the experiment, 30 × 10^3^ cells/mL of the corresponding *Microcystis* sp. strain were administered every four days. Survival was monitored on days 5, 9 and 12 of the experiment (Fig. S3). Experimental units were kept at 19 ± 1°C, 16:8 h light–dark regime.

### Statistical analyses

Throughout the following, we refer to ‘*Daphnia* timing’ as the variable indicating whether early (0) or late (1) *Daphnia* are considered, to ‘*Microcystis* timing’ as the variable indicating whether *Daphnia* are exposed to early (0) or late (1) *Microcystis* strains and to ‘contemporality’ as the variable indicating whether *Daphnia* was exposed to *Microcystis* with a different (0, allotemporal) or matching timing (1, contemporal).

As our main analysis, we use a binomial generalized linear mixed model (GLMM) with a logistic link function to model survival at the twelfth day, i.e. the final experimental day. The number of surviving individuals *y*_*i*_ out of the total number of individuals *n*_*i*_ in experimental unit *i* is modelled as follows:

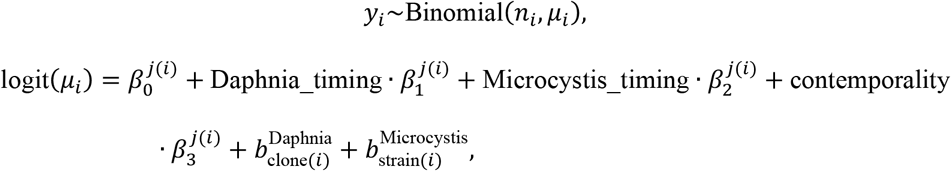

where 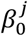is the intercept, 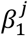is the effect of *Daphnia* timing, 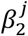is the effect of *Microcystis* timing, 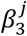is the effect of contemporality, 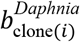 is a normally distributed random effect for the *Daphnia* clonal lineage clone(*i*)used in experimental unit *i* and 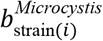 is a normally distributed random effect of the *Microcystis* strain(*i*)used in experimental unit *i*. To allow model parameters to vary across the two years, two distinct sets of regression coefficients are used (i.e. one set for 2018 and one set for 2019), with the superscript *j*(*i*) referring to the year of experimental unit *ii*. In an alternative model specification, a third random effect 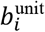 is included at the level of the experimental unit, to account for any overdispersion which might be caused by, for instance, additional genotype x genotype interactions unrelated to contemporality left unexplained by additive genotype effects captured by the *Daphnia* clonal lineage and *Microcystis* strain random effects alone.

We implemented this model using the probabilistic programming language Stan (Carpenter et al., 2017) through the ‘cmdstanr’ package in R v.4.3.1 (R Core Team, 2023). Stan performs Bayesian inference by means of a dynamic Hamiltonian Monte Carlo (HMC) algorithm, a gradient-based Markov chain Monte Carlo (MCMC) sampler (Carpenter et al., 2017). We used zero-centred, weakly informative priors with scale 3 for the variables *β*_0_, *β*_1_, *β*_2_and *β*_3_. Similarly, we used weakly informative half-normal priors with scale 3 for the scales of the random clonal lineage and strain effects 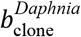 and 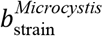. We ran eight chains with 2,000 iterations each, of which the first 1,000 were discarded as warm-up. Model convergence was confirmed both visually by means of traceplots and numerically by means of effective sample sizes, the absence of divergent transitions and the Potential Scale Reduction Factor, for which all parameters had 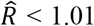 (Vehtari et al., 2021). A satisfactory goodness-of-fit was confirmed through posterior predictive checks (Gelman et al., 1996). We performed prior sensitivity analyses to confirm that model results are robust against alternative prior specifications. Traceplots and visualizations of the posterior predictive checks and prior sensitivity analyses are provided in the Supplementary Information (Fig. S4-9). We used the ‘tidybayes’ v.2.3.1 package (Kay, 2022) to visualize the posterior distributions of the inferred effects and their corresponding derived quantities.

To conform readers who are more familiar with frequentist statistics, we also fitted the binomial GLMM using maximum likelihood with the ‘glmer’ function of the ‘lme4’ package v.1.1 (Bates et al., 2015) in R v.4.3.1 (R Core Team, 2023). Rather than estimating two sets of parameters for both years within the same model (which is not readily possible using the considered software) or including interactions between all regression parameters and the year, we estimate two separate models in a year-by-year fashion.

To ensure that our results are insensitive to the choice of survival at day 12 as endpoint, we also performed a Bayesian interval-censored frailty survival analysis (Bogaerts et al., 2017) to model individual survival over the entire experiment′s duration. We outline methodological details for this analysis in the Supplementary Information (Section S1).

## Results

*Daphnia* survival after twelve days of exposure to *Microcystis* varied widely across experimental units (Fig. 1; Fig. S3). The average observed survival rate across individuals was 37.2%.

**Figure 1.**
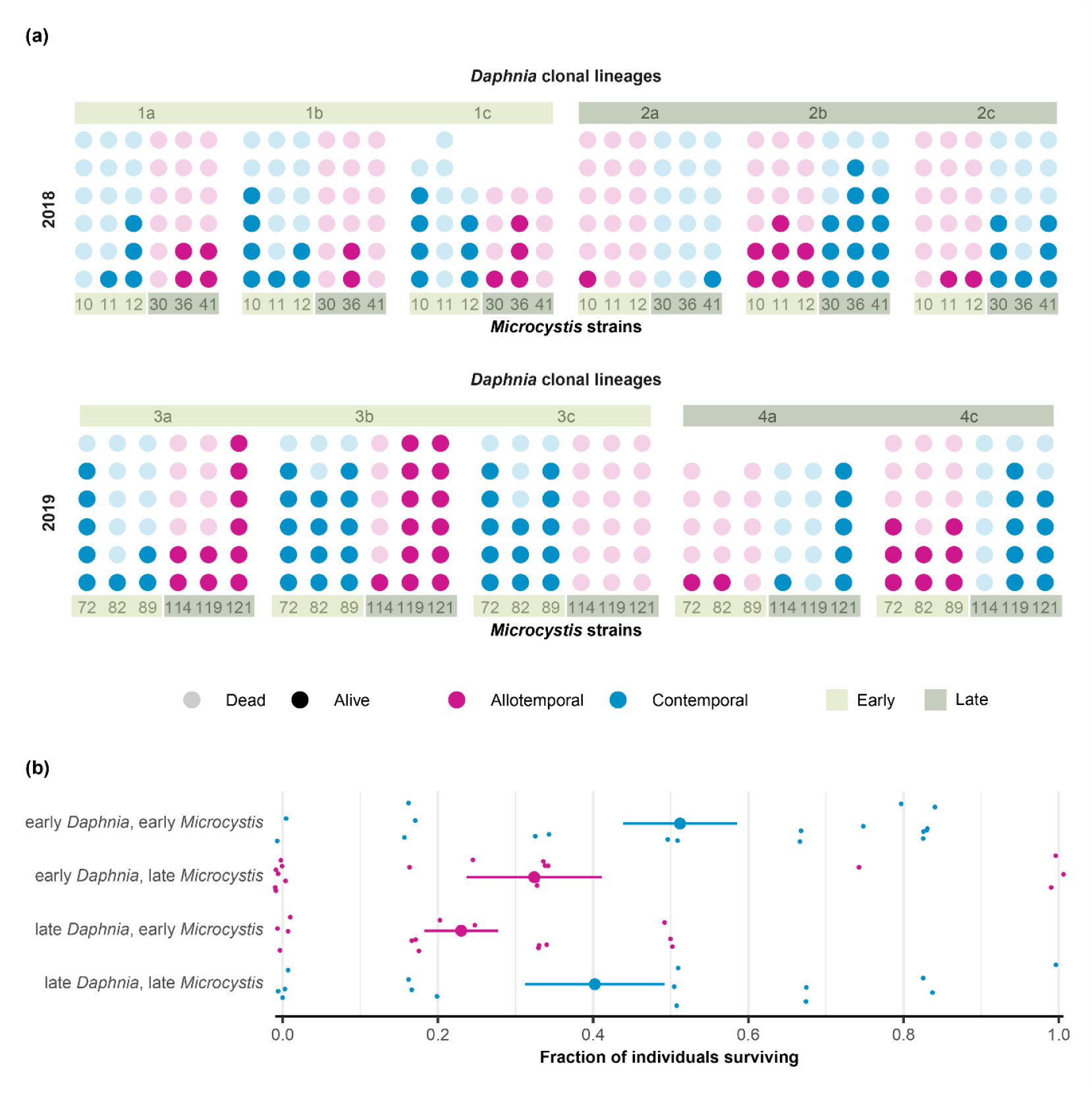
Observed *Daphnia* survival at day 12. **(a)** Survival status of each individual, represented by a point (dead: transparent; alive: opaque), organized per *Daphnia* clonal lineage and per *Microcystis* strain. Allotemporal combinations are shown in pink and contemporal combinations are shown in blue. **(b)** Averaged survival for each combination of *Daphnia* clonal lineage and *Microcystis* strain. Grand means and standard errors for each combination of early and late *Daphnia* and *Microcystis* clonal lineage are shown by means of big dots and horizontal lines. Allotemporal combinations are shown in pink and contemporal combinations are shown in blue, jittered horizontally and vertically to a small extent to avoid overlap.

**Figure 2.**
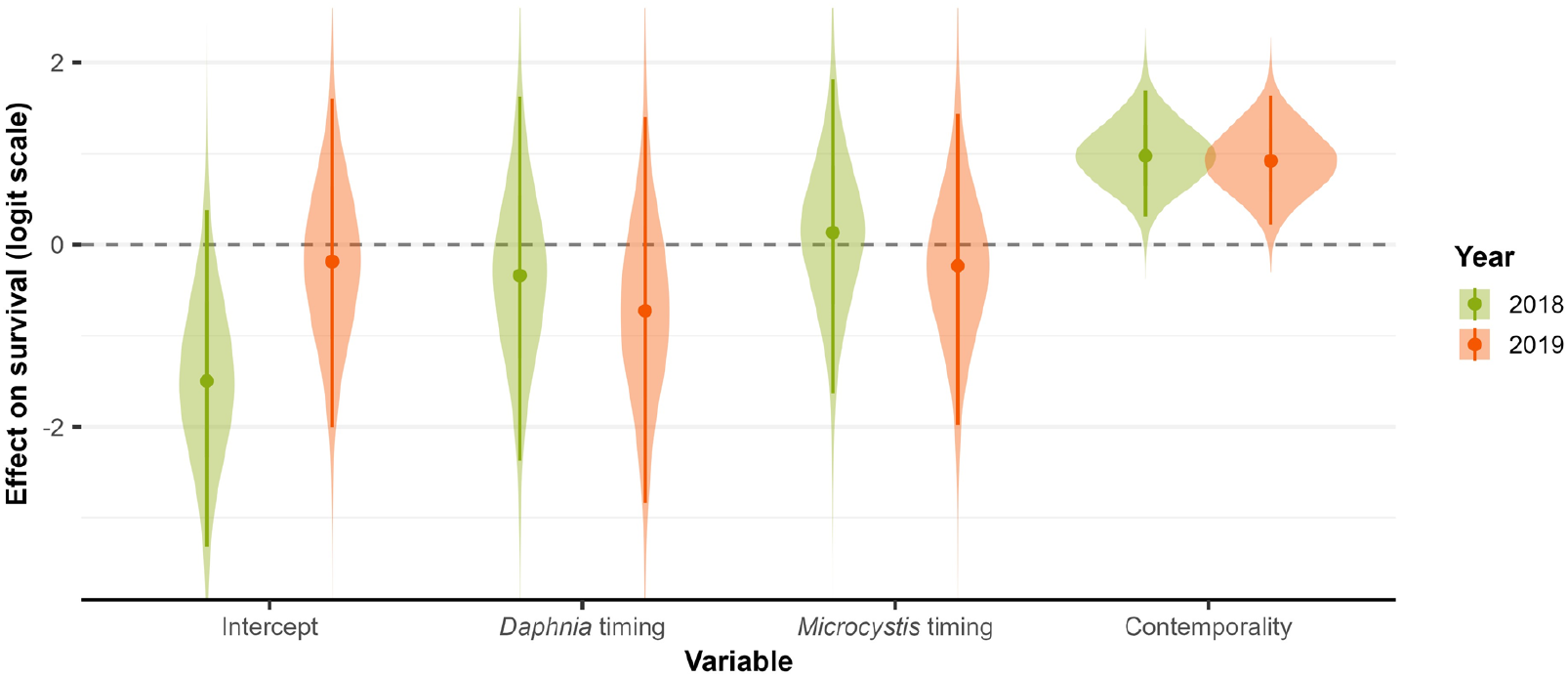
Posterior distributions of the model intercept and the effects of *Daphnia* timing, *Microcystis* timing and contemporality on *Daphnia* survival (logit scale) at the twelfth day, for 2018 (green) and 2019 (orange), as obtained from the binomial GLMM. The posterior medians and 95% credible intervals are represented by dots and vertical lines respectively.

Using a binomial GLMM to analyse survival at day 12, we found no evidence for differential survival between early and late *Daphnia* clonal lineages (posterior probability of a positive difference, averaged across both years: 70.9%), neither did we find evidence for a difference between early and late *Microcystis* strains (av. post. prob.: 52.4%). We did, however, find very strong statistical support for a positive effect of contemporality (av. post. prob.: 99.6%), with contemporal combinations featuring a higher survival compared to allotemporal ones (Figs. 1-2). Averaged across both years, the posterior mean temporality effect equals 0.95 (95% CrI [0.27,1.67]) on the logit scale. This corresponds to a 17.7 % decrease in posterior mean survival probability when *Daphnia* are exposed to *Microcystis* strains from another moment (Fig. S10). Moreover, these patterns are strongly replicated across both years, with only 54.7% posterior probability of a positive or negative difference in the effect of contemporality between both years. For reference, a year-by-year frequentist binomial GLMM analysis yields highly similar results, with p-values of 0.006 and 0.013 for the effect of contemporality in 2018 and 2019, respectively (Table S2).

The interval-censored frailty survival analysis, in which data of all time points is integrated, yields qualitatively identical results as the binomial GLMM analysis performed on the twelfth day (Fig. S11). The survival functions generated through this interval-censored frailty survival analysis visually confirm that contemporal combinations host an increased survival compared to allotemporal combinations, while there is much more overlap in confidence intervals among survival curves of early and late *Daphnia* or of *Daphnia* exposed to early and late *Microcystis* (Fig. S12).

Posterior distributions for the different *Daphnia* genotypes and *Microcystis* strains show that there is much variation in survival upon exposure to *Microcystis* among *Daphnia* genotypes (Figs. 3-4). Similarly, there is variation in survival of *Daphnia* when exposed to different *Microcystis* strains, reflecting that *Microcystis* genotypes differ in toxicity (Figs. 3-4). For instance, *Daphnia* clonal lineages 2a and 2b feature the highest intrinsic difference in tolerance (Figs. 3-4; posterior probability of a difference: > 99.9%). On the other hand, *Daphnia* individuals exposed to *Microcystis* strain 114 have a distinctly lower survival probability at day 12 than individuals exposed to *Microcystis* strain 121 (posterior probability of a difference: > 99.9%).

**Figure 3.**
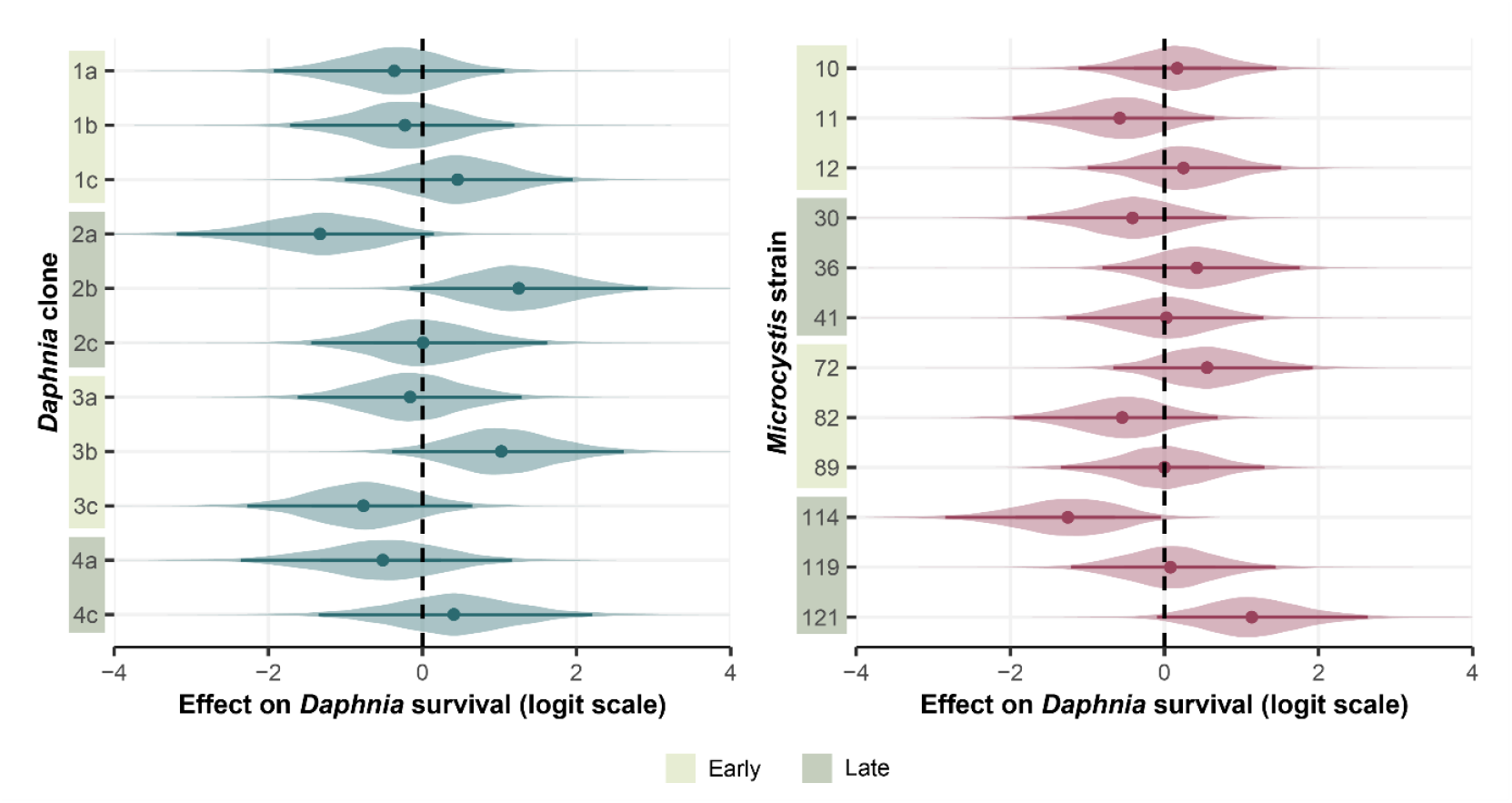
Posterior distributions of the *Daphnia* clonal lineage (left) and *Microcystis* strain (right) effects on *Daphnia* survival (logit scale) at the twelfth day, as obtained from the binomial GLMM. These effects represent the clone- and strain-specific variation, corrected for the effects of year, *Daphnia* timing, *Microcystis* timing, contemporality, *Microcystis* strain (for the *Daphnia* clonal lineage effects only) and *Daphnia* clone (for the *Microcystis* strain effects only). The posterior medians and 95% credible intervals are represented by dots and horizontal lines respectively.

**Figure 4.**
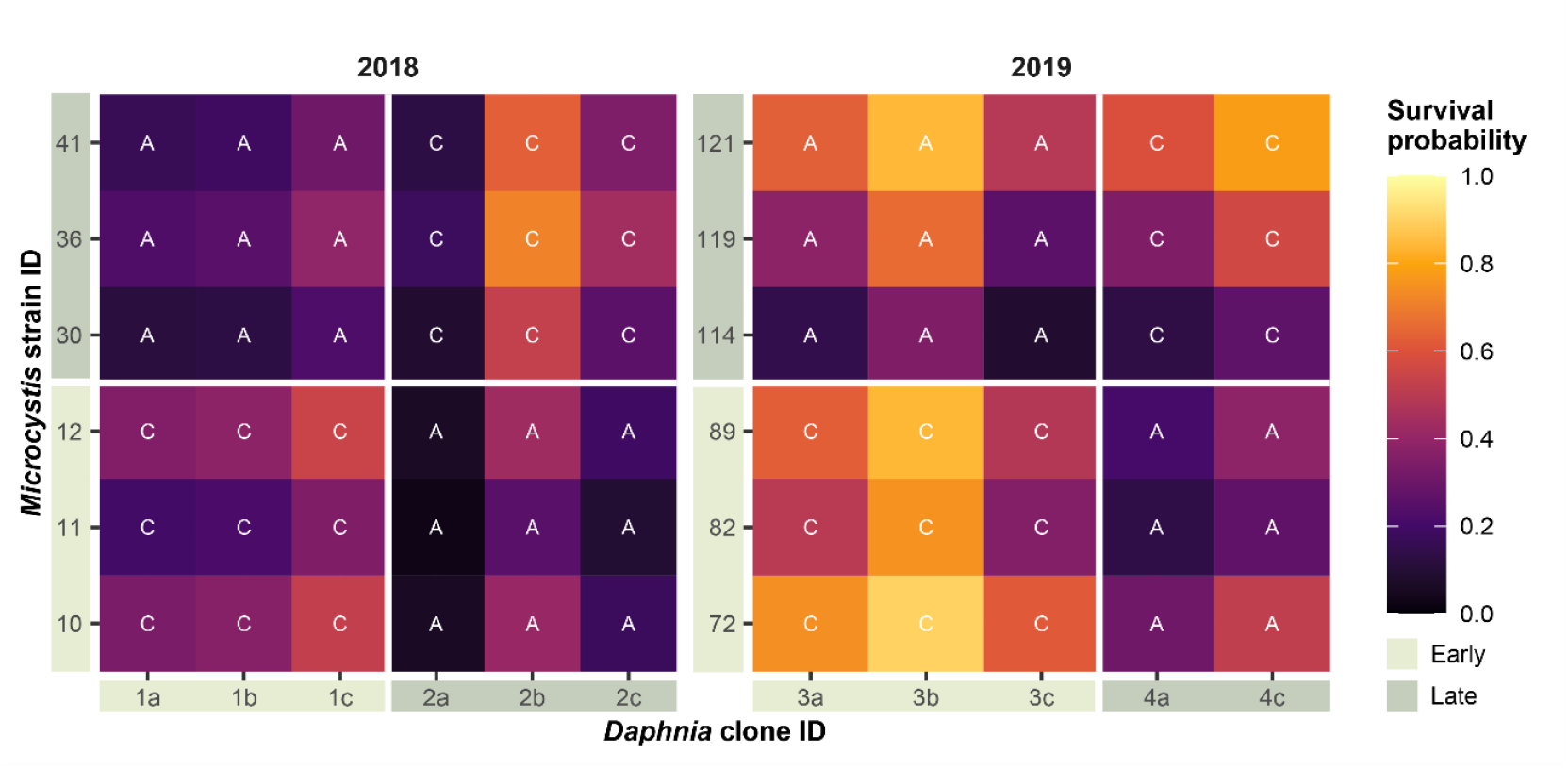
Heatmap showing the posterior mean modelled survival probability for each combination of *Daphnia* clonal lineages and *Microcystis* strains, as obtained from the binomial GLMM. White letters indicate whether the combination of *Daphnia* clonal lineage and *Microcystis* strain pertains to a contemporal (‘c’) or allotemporal (‘a’) condition.

A comparison of the main binomial GLMM with a fully saturated binomial GLMM, in which an observation-level random effect (i.e. a *Daphnia* genotype x *Microcystis* genotype random effect) is added, indicates that our main model can explain a posterior mean fraction of 67.3% (95% CrI [53.4,79.2]) of the saturated linear predictor’s variation (Fig. S13). Hence, approximately two thirds of the variation in survival can be explained by contemporality, *Daphnia* genotype and *Microcystis* genotype alone. Approximately one third of the variation is left unexplained, suggesting that genotype x genotype effects unrelated to contemporality, or other sources of overdispersion, were also important in determining *Daphnia* survival in our experiment. While our experimental design allows to disentangle additive effects of *Daphnia* genotype and *Microcystis* genotype as well as the genotype x genotype interaction induced by contemporality, variation induced by additional genotype x genotype interactions might be confounded with observational noise because of the absence of replicates at the level of *Daphnia* genotype x *Microcystis* genotype combination.

## Discussion

In this study, we exposed early and late season genotypes isolated from a natural *D. magna* population over two years to *Microcystis* strains isolated from the same two time periods of their respective year of isolation. Our results do not only show that *Daphnia* genotypes differ in sensitivity to *Microcystis* and *Microcystis* strains differ in toxicity to *Daphnia*, but also reveal a striking pattern of higher survival when *Daphnia* genotypes are exposed to *Microcystis* strains that were isolated during the same time period compared to *Daphnia* that were exposed to *Microcystis* strains that were isolated during a different period of the growing season.

*Daphnia* populations have been shown to be genetically differentiated in their capacity to cope with *Microcystis* (Chislock et al., 2019; Sarnelle & Wilson, 2005; Schwarzenberger et al., 2020) and that *Daphnia* can become more tolerant to *Microcystis* through acclimatization (Chislock et al., 2019; Sarnelle & Wilson, 2005), epigenetic effects (Asselman et al., 2017; Ger et al., 2016) or through microbiome-mediated acclimatization (Macke et al., 2017). Influential studies have shown that *Daphnia* can genetically adapt to the presence of *Microcysti*s over the course of decades in resurrection ecology studies (Hairston et al., 1999; Isanta-Navarro et al., 2021; Jiang et al., 2016). Our results suggest that genetic adaptation of *Daphnia* to *Microcystis* can occur over the course of one growing season and enables *Daphnia* to effectively cope with contemporal *Microcystis* populations. While adapting to changes in the *Microcystis* population, *Daphnia* lose some of their tolerance to past populations of *Microcystis*. This is reflected by the fact that late season *Daphnia* populations suffer from exposure to the early-season *Microcystis* strains. This rapid adaptation is expected to influence seasonal phytoplankton and *Microcystis* dynamics and might explain the genetic adaptation affecting *D. mendotae* seasonality and phytoplankton biomass in Oneida Lake, as observed by Schaffner et al. (2019). The PEG model explaining seasonal changes in phytoplankton and zooplankton biomass in lakes might in part be driven by the eco-evolutionary interactions revealed for a deep lake by Schaffner at al. (2019) and the shallow pond in our study.

We show strong *Daphnia* genotype x *Microcystis* genotype interactions. These G x G interactions are a prerequisite for the pattern of contemporal adaptation that we observe, as it allows the population of the one interacting species to not only adapt to *Microcystis* as a stressor, but to adapt to the specific genotype composition of *Microcystis. Microcystis* strains are known to strongly differ in their toxicity, both in terms of the capacity to produce microcystins and the amount they produce, but also in terms of the synthesis of other polypeptides with potentially toxic effects (Xu et al., 2023) or secondary metabolites such as digestive protease inhibitors (Schwarzenberger et al., 2020). In addition to variation in toxicity, some *Microcystis* strains also exhibit inducible colony formation in response to grazing, impeding ingestion. Considering this wide variety of toxicity profiles and defense strategies, our results suggest that *Daphnia* genotypes are differentially able to adapt to different *Microcystis* strains, enabling clonal selection. Lemaire et al. (2011) also showed important genotype x genotype interactions in determining the performance of *Daphnia* when exposed to *Microcystis*. They used *Daphnia* clonal lineages and *Microcystis* strains that were isolated from habitats that were geographically isolated, including *Daphnia* from both the European and African continent. The present study shows that the same pattern exists within a single habitat, and contributes to our understanding of seasonality of zooplankton-phytoplankton interactions. Our results also align with the seasonal dynamics in the clonal composition of *D. magna* observed in the field by Schwarzenberger et al. (2013), which might be driven by corresponding dynamics in the cyanobacterial composition and the protease inhibitors they produce (Schwarzenberger et al., 2013). The results of our experiments demonstrate that *Daphnia* is indeed able to seasonally track changes in the composition of cyanobacterial strains.

Our results testify to the large intrapopulation genetic variation that is regularly observed in *Daphnia* (Chaturvedi et al., 2021; De Coninck et al., 2013; Ilić et al., 2021) and that can explain why local adaptation has been reported in *Daphnia* populations in response to a wide range of environmental gradients and change such as temperature (Brans & De Meester, 2018; Geerts et al., 2015), pollution (Almeida et al., 2023; Cuenca-Cambronero et al., 2021; De Coninck et al., 2013), UV light (Miner & Kerr, 2011), including interactions with antagonists such as predation (Stoks et al., 2015), parasites (Decaestecker et al., 2007) and competitors (Steiner, 2004). *Daphnia-Microcystis* interactions are especially interesting because both interactors can kill each other if they are sufficiently adapted to their enemy. This may result in fast and reciprocal responses, similar to those reported for host-parasite interactions (Decaestecker et al., 2007, 2013; Gandon et al., 2008). Similar to the strong genotype x genotype interactions that are often observed in host-parasite coevolution (Capaul & Ebert, 2003; Ebert, 2008), we also observe strong *Daphnia* genotype x *Microcystis* genotype interactions, both at larger spatial scales (Lemaire et al., 2011) as well as within single habitats (present paper).

The strong *Daphnia* genotype x *Microcystis* genotype interactions might not only enable *Daphnia* to adapt to *Microcystis* but may also allow *Microcystis* populations to adapt to the changes in the *Daphnia* population, leading to continuous dynamics of coevolution (Anderson & May, 1982; Ehrlich & Raven, 1964; Ley et al., 2008). Our results show that *Daphnia* populations seem to be able to track changes in *Microcystis* populations during a growing season (i.e. late-season *Daphnia* genotypes are better adapted to late-season *Microcysti*s than early-season *Daphnia* genotypes) and show that this adaptation has a cost in terms of susceptibility to early-season *Microcystis* (i.e. late-season *Daphnia* genotypes are less well adapted to early-season *Microcysti*s than early-season *Daphnia* genotypes). The fact that early-season *Daphnia* are more susceptible to late-season *Microcystis* might reflect adaptation of the *Microcystis* population to *Daphnia*, driven by selection against grazing. Given that we did not carry out genomic analyses on the different *Microcystis* strains used in this experiment, we cannot be certain that evolution is involved rather than epigenetic changes or a shift in relative abundances of closely related *Microcystis* species. Recently, it has been shown that also the microbiome can structure such G x G interactions resulting in G x M x G interactions and modified phenotypes with increased *Microcystis* tolerance (Macke et al., 2017). If, however, both *Daphnia* and *Microcystis* evolve in response to each other, the *Daphnia-Microcystis* interaction might be a nice example of a geographic mosaic of coevolution in which temporal adaptation result in spatial variation of coevolutionary hotspots and coldspots (Thompson, 2005). Modified locally adapted *Daphnia* microbiomes have been shown to mediate tolerance to *Microcystis* (Houwenhuyse et al., 2021). These aspects have high potential relevance with respect to the management of *Microcystis* blooms.

Overall, our study demonstrates the importance of short-term eco-(co)evolutionary dynamics in determining interactions between cyanobacteria and zooplankton grazers. We show that the composition of cyanobacteria and their toxicity for zooplankton grazers varies over short periods of time, and that the zooplankton grazer studied, the water flea *D. magna*, is able to adapt to these changes in order to reduce mortality imposed by *Microcystis*. This mechanism likely contributes to the ability of *Daphnia* populations to delay or even prevent cyanobacteria blooms in freshwater ecosystems (Benndorf & Henning, 1989; de Bernardi & Guissani, 1990; Jeppesen et al., 1996; Matveev et al., 1994; Søndergaard et al., 2008; Vanni, 1984). It may also explain why the literature on the toxicity of *Microcystis* and the capacity of *Daphnia* to graze on and control *Microcystis* is so divergent in its conclusions, as the outcome may depend on who of the two interacting partners in the particular setting studied is at the moment of sampling winning in the coadaptation, and because most studies combine *Microcystis* and *Daphnia* strains that do not co-occur. The eco-evolutionary dynamics that may result from the dynamics reported in our study might impact the observed seasonality and food-web interactions in lakes and ponds (Schaffner et al., 2019).

## Data and code availability statement

The data and full code for the analysis is available through the following GitHub repository: https://github.com/MFajgenblat/Daphnia_Microcystis_temporal_dynamics.

## Author contributions

AB, LDM and ED conceived the study. AB, KC, EvdB and IV performed the experiment. MF and MC lead the statistical analysis of the data. MF, EG and LDM lead the writing of the manuscript. RA provided critical feedback during the writing process. All mentioned co-authors contributed substantially to the further refinements of the manuscript.

## Acknowledgements

We gratefully acknowledge the NGO Vrienden van Heverleebos en Meerdaalwoud and its volunteers for providing access to Langerodevijver and granting permission for sampling, which was essential for this study. We thank Laura Herregodts and Cedric De Witte-Vroman for their valuable practical help in performing this experiment.

