## Supplementary Information for "Rapid temporal adaptation structures tolerance to toxic cyanobacteria in a natural population of the water flea *Daphnia*"

#### Extended methods

##### *S1 Bayesian interval-censored frailty survival analysis*

We use an interval-censored frailty survival analysis approach to model individual survival over the entire experiment's duration. We assume an exponential survival function with a rate $\lambda_i$ that varies across experimental units:

$$S_i(t) = e^{-\lambda_i t},$$

where $t$ is the time since the experiment was initiated, expressed in the number of days divided by 10. The survival rate $\lambda_i$ of individual $i$ is modelled in an analogous fashion as the survival probabilities in the binomial GLMM:

$$\log(\lambda_i) = \beta_0^{j(i)} + \text{Daphnia\_timing} \cdot \beta_1^{j(i)} + \text{Microcystis\_timing} \cdot \beta_2^{j(i)} + \text{contemporality} \cdot \beta_3^{j(i)} + b_{\text{clone}(i)}^{\text{Daphnia}} + b_{\text{strain}(i)}^{\text{Microcystis}}.$$

Since survival is only assessed at $t = 0,0.5,0.9,1.2$, the survival time of each individual $i$ is only available at a resolution of the four time intervals $[0,0.5)$, $[0.5,0.9)$, $[0.9,1.2)$ and $[1.2,+\infty)$, each being defined by the lower interval bound $l_i$ and the upper interval bound $u_i$. Hence, the joint likelihood for all individuals $i = 1,2,\dots,n$ is defined as:

$$L(\boldsymbol{\beta}_0, \boldsymbol{\beta}_1, \boldsymbol{\beta}_2, \boldsymbol{\beta}_3, \boldsymbol{b}^{Daphnia}, \boldsymbol{b}^{Microcystis} | \boldsymbol{l}, \boldsymbol{u}) = \prod_i^n \left(e^{-\lambda_i u_i} - e^{-\lambda_i l_i}\right).$$

We implemented this model in Stan, following the same approach and using the same prior specifications as for the binomial GLMM presented in the main text.

### Supporting figures

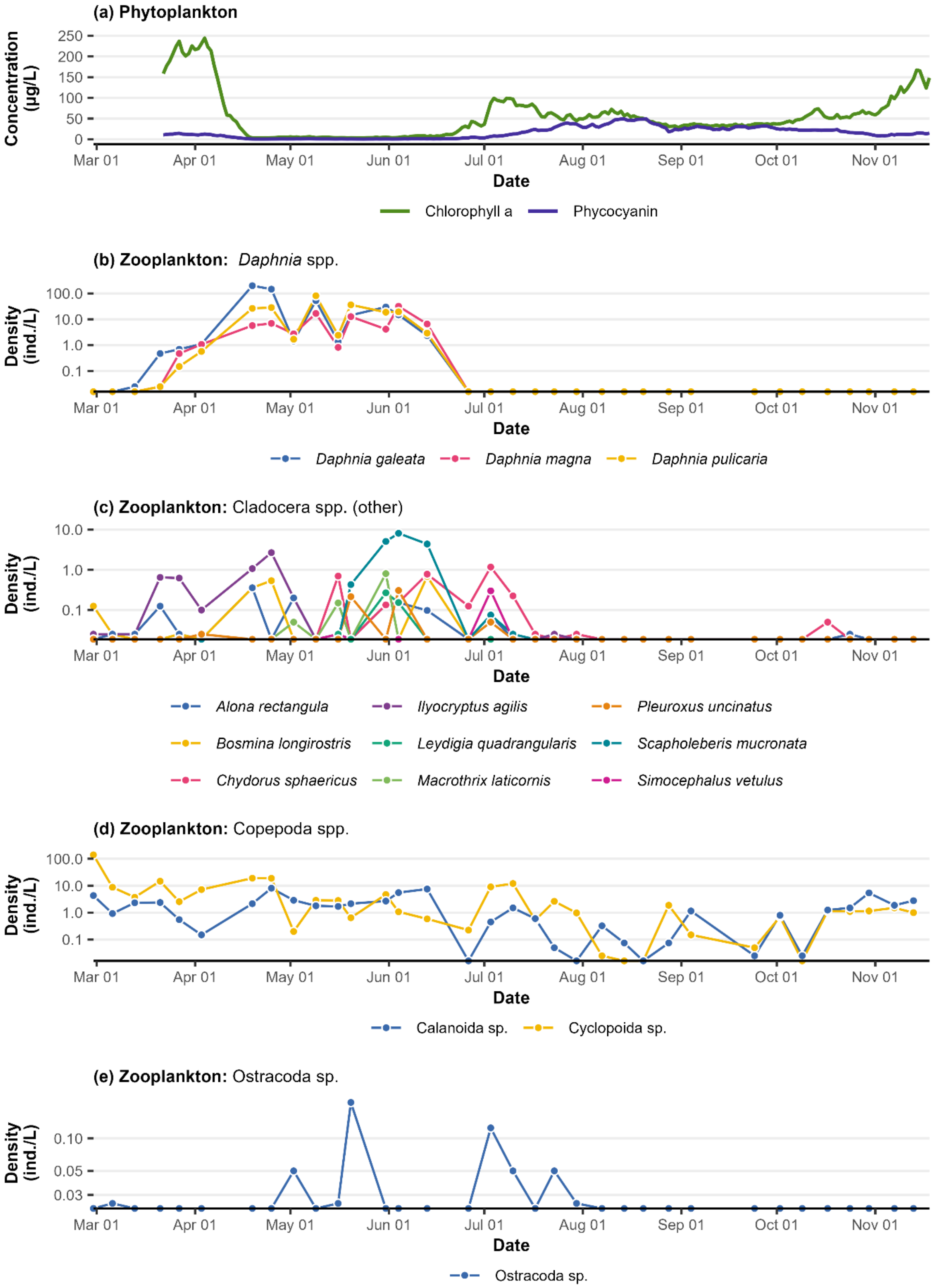

**Figure S1.** Temporal dynamics of phytoplankton and zooplankton in Langerodevijver from February to November 2019. **(a)** Daily median *in vivo* concentrations of chlorophyll a and phycocyanin, from hourly measurements with two EXO2 sondes (YSI Incorporated, Yellow Spring, USA) placed in Langerodevijver. **(b-e)** Observed densities of zooplankton taxa, by identification of specimens in depth integrating water samples of 30 L, assessed on a weekly basis (following the protocol outlined in (De Bie et al., 2012)). The y-axis (density of individuals) is scaled logarithmically for clarity.

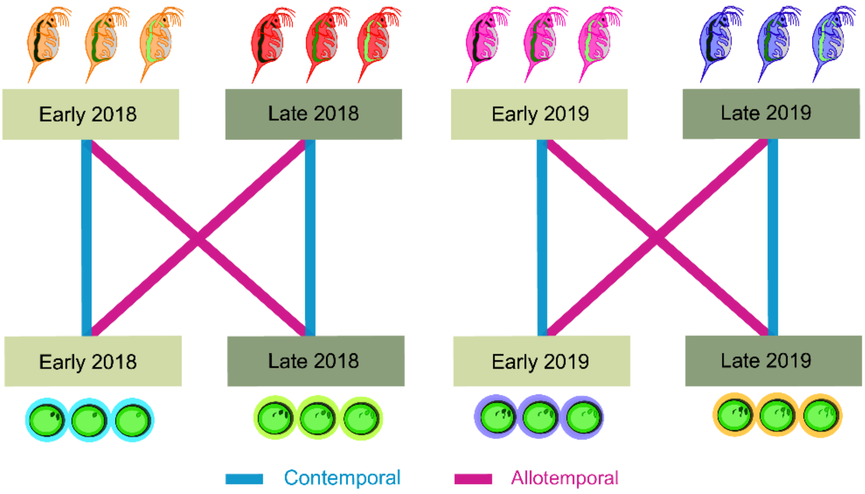


**Figure S2**. Graphical representation of the experimental design.

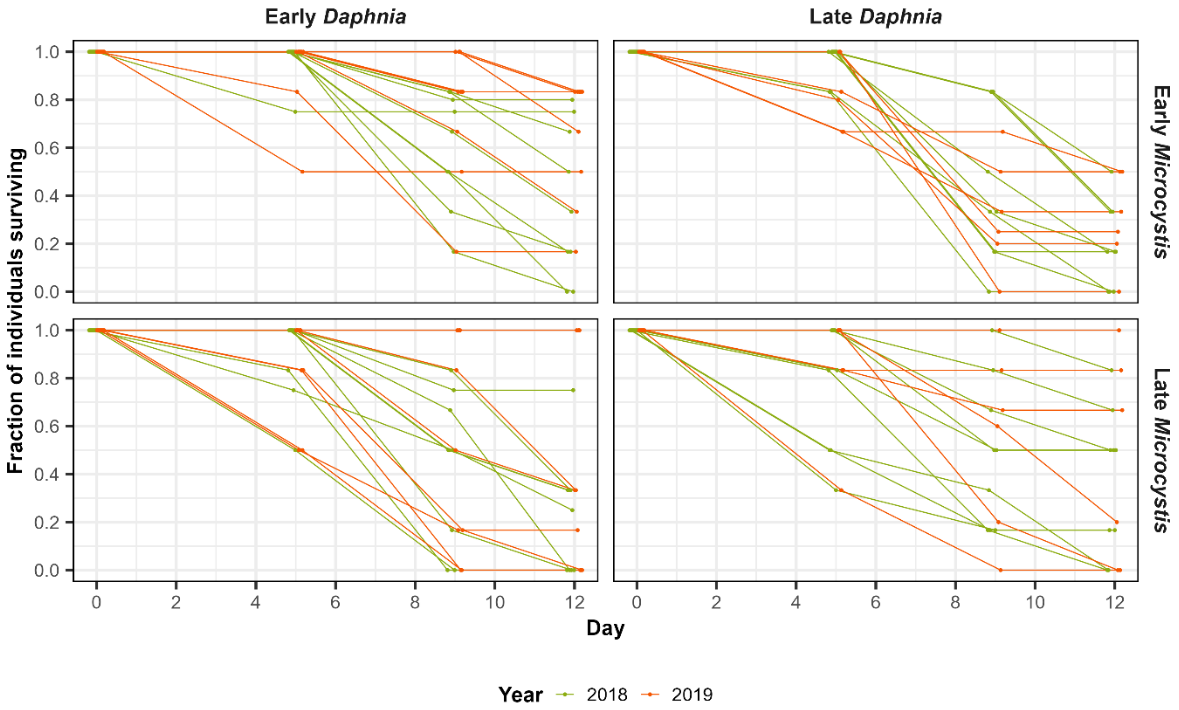


**Figure S3.** Visualization of the raw survival data. For each experimental unit, the observed survival percentage at days 0, 5, 9 and 12 is shown by means of a dot. Dots belonging to the same experimental units are joined by a line, and both dots and lines are dodged horizontally by a small extent to avoid overlapping. The two experimental years are represented by different colours.

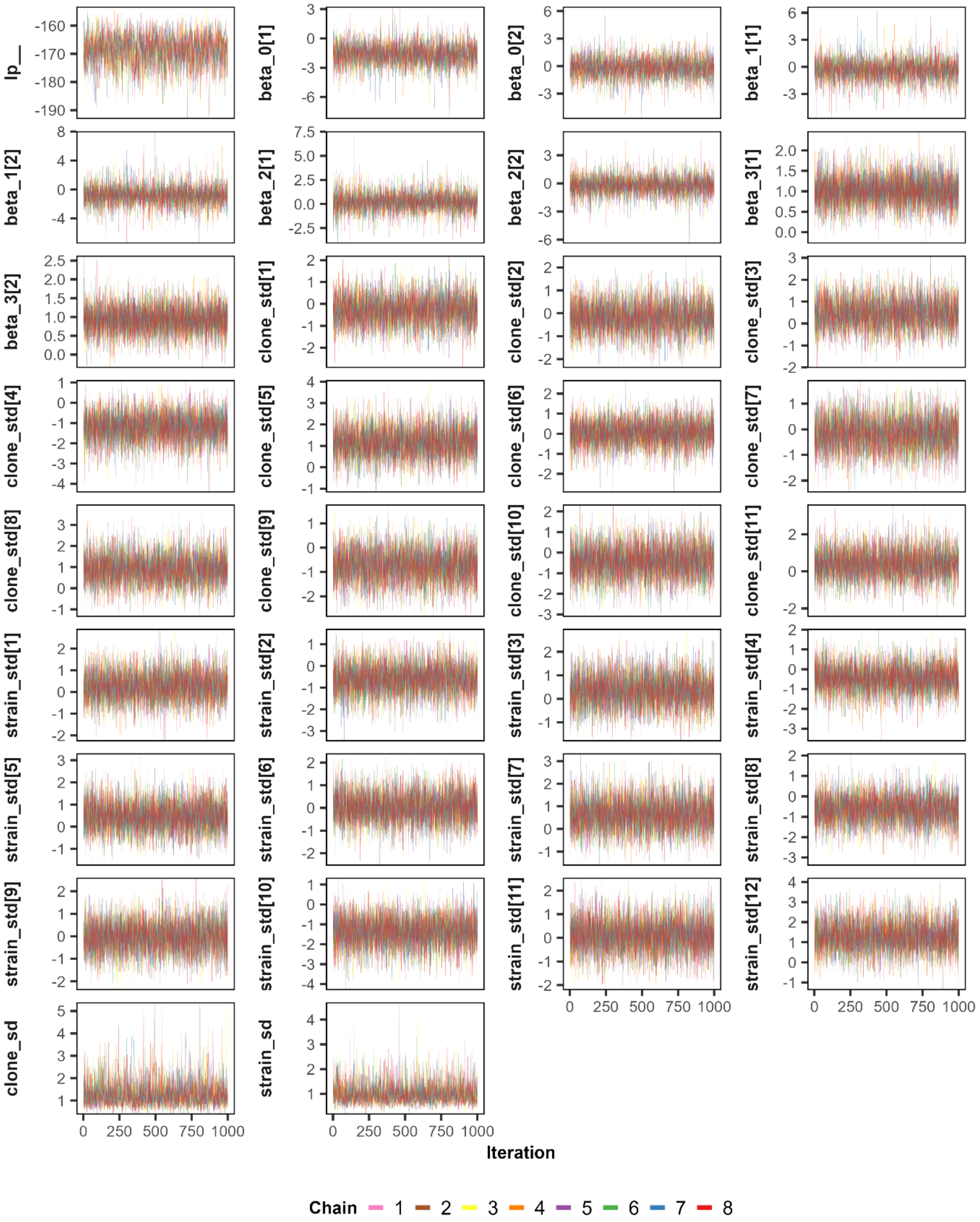


**Figure S4.** Traceplots for the parameters of the binomial GLMM analysis. Each MCMC chain is shown by a different colour. Warmup draws are not shown.

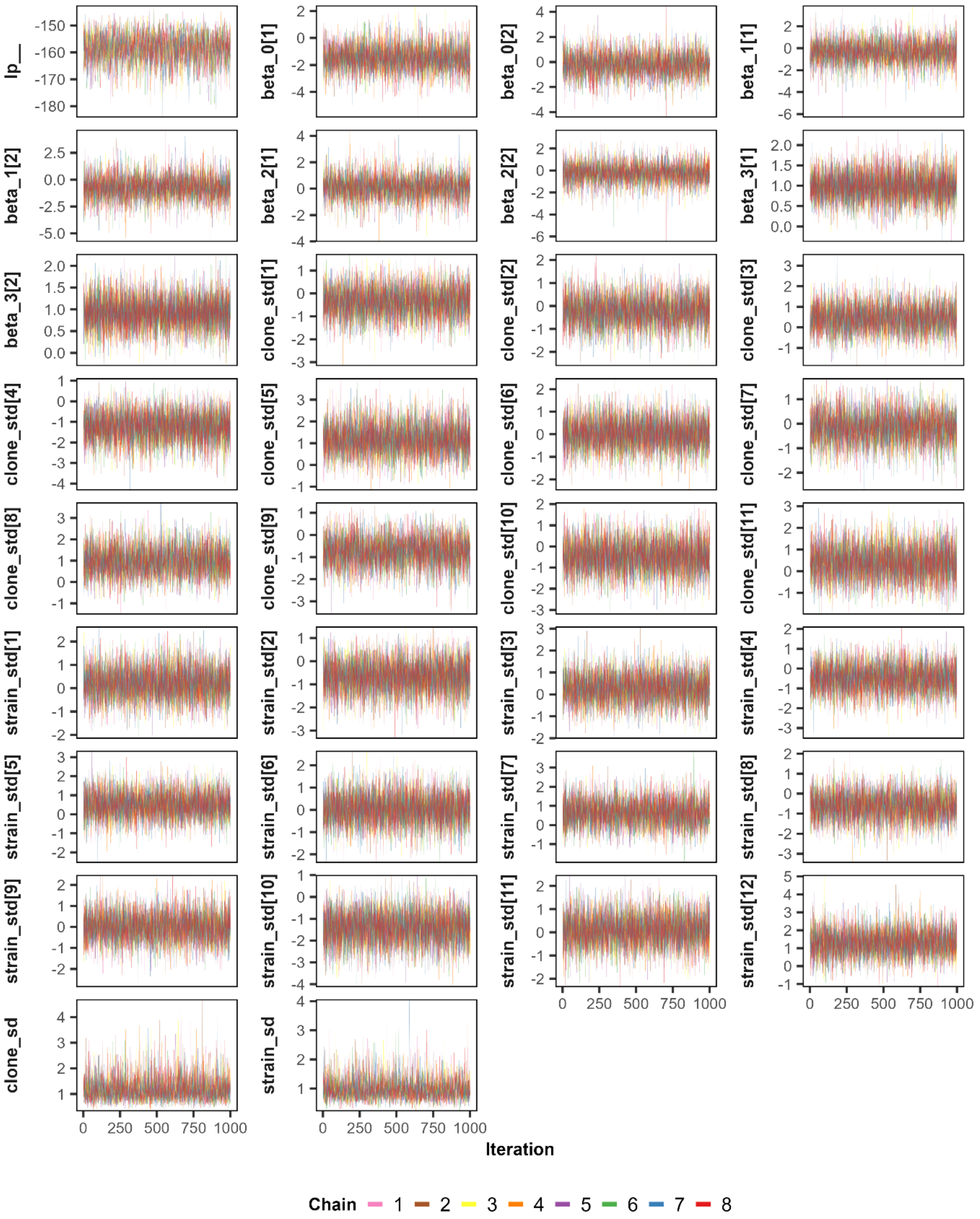


**Figure S5.** Traceplots for the parameters of the interval-censored frailty survival analysis. Each MCMC chain is shown by a different colour. Warmup draws are not shown.

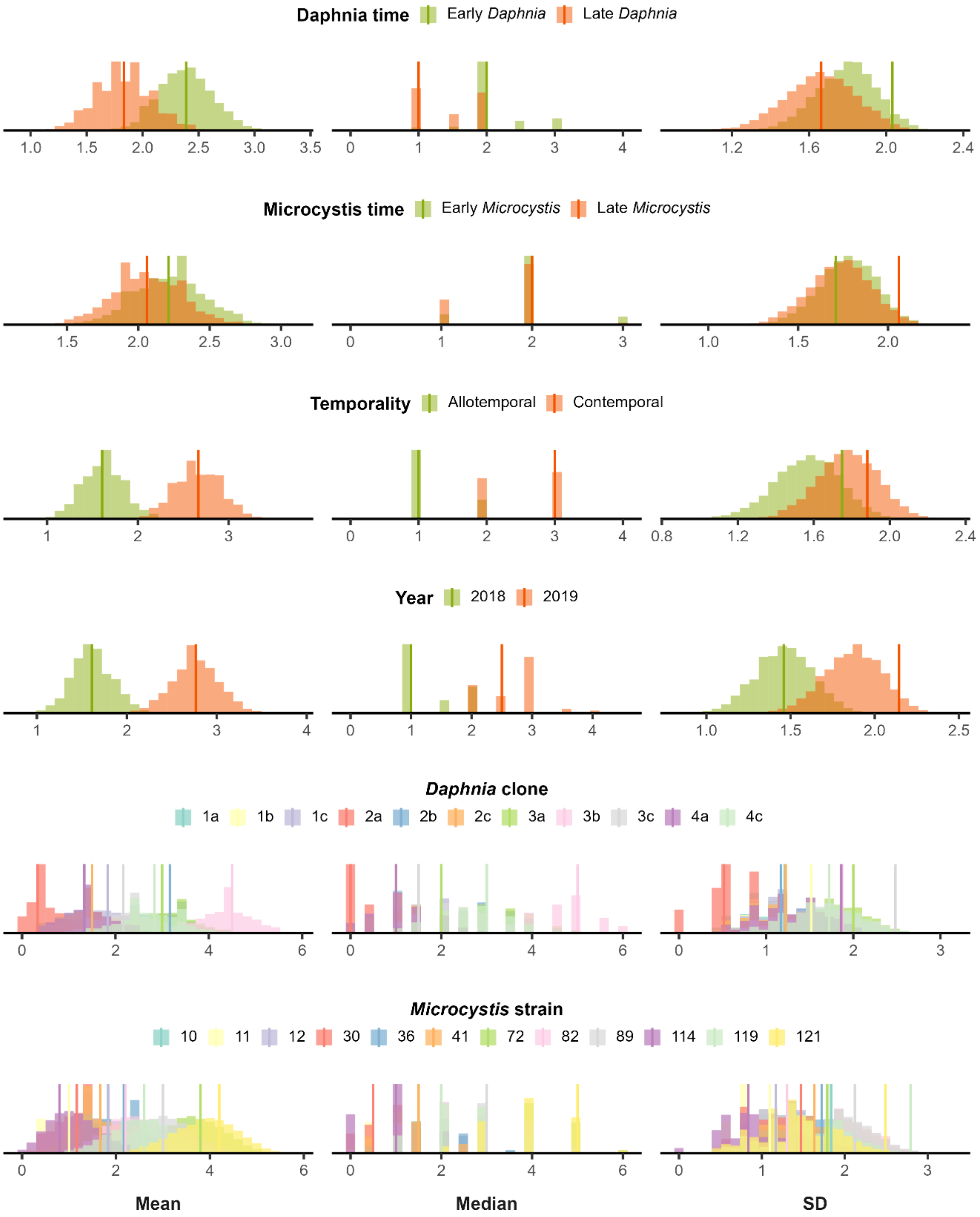


**Figure S6.** Posterior predictive checks for the binomial GLMM analysis. Posterior predictive densities are shown by means of histograms, coloured by various subgroups (vertically stacked subplots). The three columns show the posterior distribution of the mean, median and standard deviation of the fraction of survived individuals across all experimental units belonging to the same group, while vertical lines show the corresponding observed mean, median and standard deviation for each group.

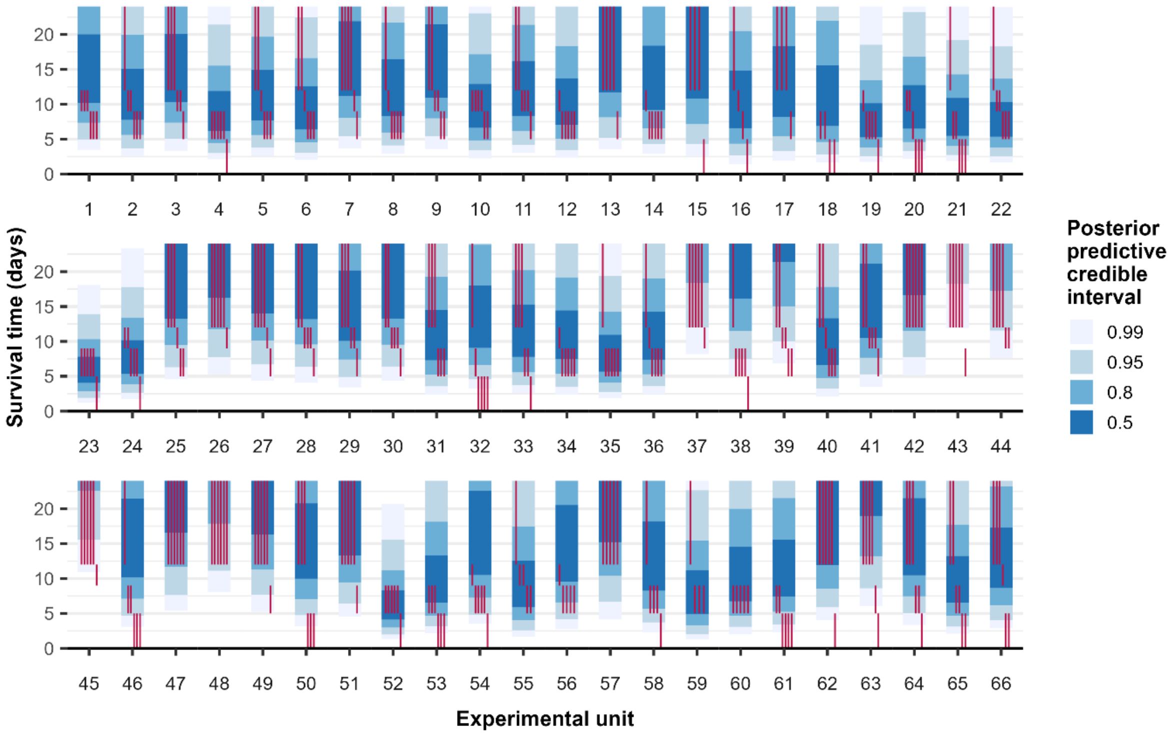


**Figure S7.** Posterior predictive checks for the interval-censored frailty survival analysis. Posterior predictive survival densities are shown by means of blue credible intervals, at uncertainty levels of 50, 80, 95 and 99%. For each experimental unit, the observed intervals during which individuals died are shown by pink vertical segments. The y-axis is truncated at day 24 for compactness.

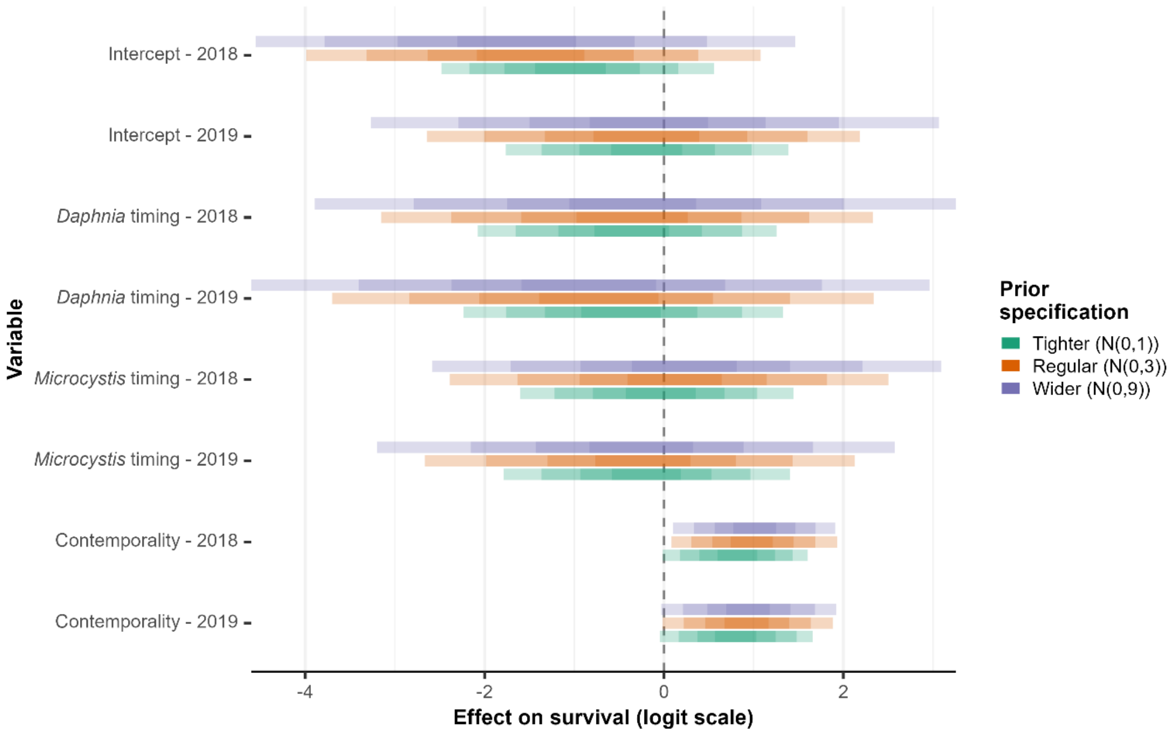


**Figure S8.** Prior sensitivity analysis for the binomial GLMM analysis, where both a tighter and wider prior specification is considered, along with the regular prior specification used in the main analysis. The tighter prior specification features Normal(0,1) and Half-Normal(0,1) priors for regression coefficients and random effect scales respectively. The wider prior specification features Normal(0,9) and Half-Normal(0,9) priors for regression coefficients and random effect scales respectively. For each prior specification, the obtained posterior distributions, as summarized by 50, 80, 95 and 99% credible intervals (with increasing transparency), of key model parameters is shown using different colours.

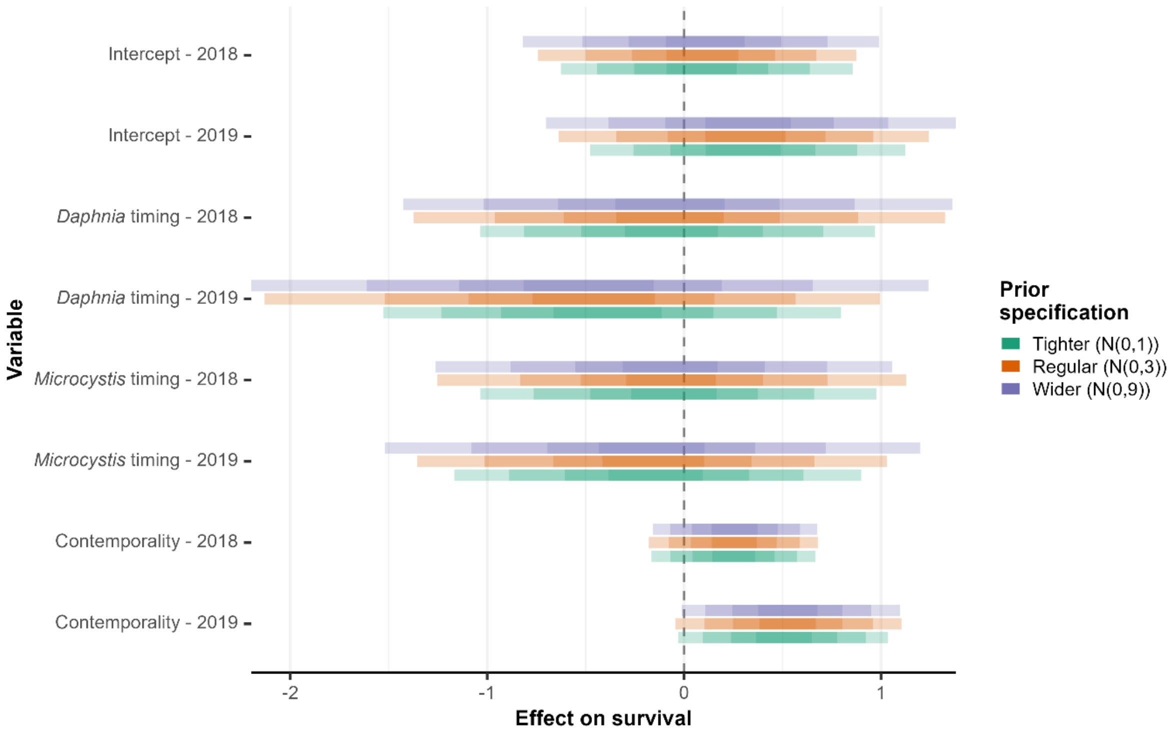


**Figure S9.** Prior sensitivity analysis for the interval-censored frailty survival analysis, where both a tighter and wider prior specification is considered, along with the regular prior specification used in the main analysis. The tighter prior specification features Normal(0,1) and Half-Normal(0,1) priors for regression coefficients and random effect scales respectively. The wider prior specification features Normal(0,9) and Half-Normal(0,9) priors for regression coefficients and random effect scales respectively. For each prior specification, the obtained posterior distributions, as summarized by 50, 80, 95 and 99% credible intervals (with increasing transparency), of key model parameters is shown using different colours.

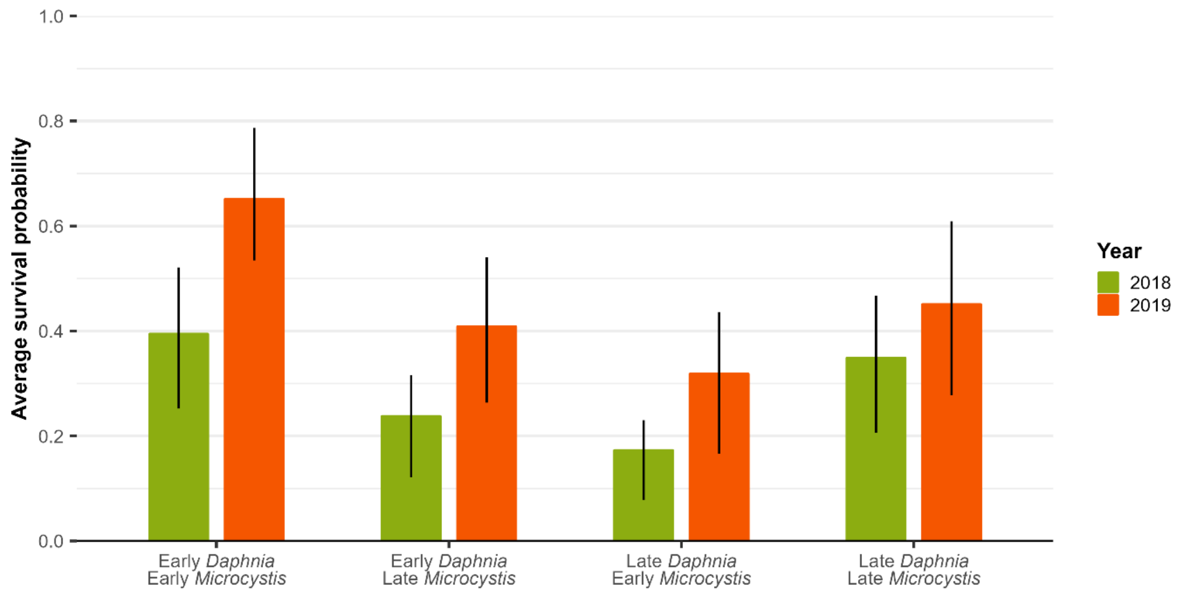


**Figure S10.** Estimated average survival (natural scale) for each combination of *Daphnia* timing and *Microcystis* timing, for 2018 and 2019, as obtained from the binomial GLMM. Posterior means and 50% credible intervals are shown by coloured bars and black vertical lines respectively.

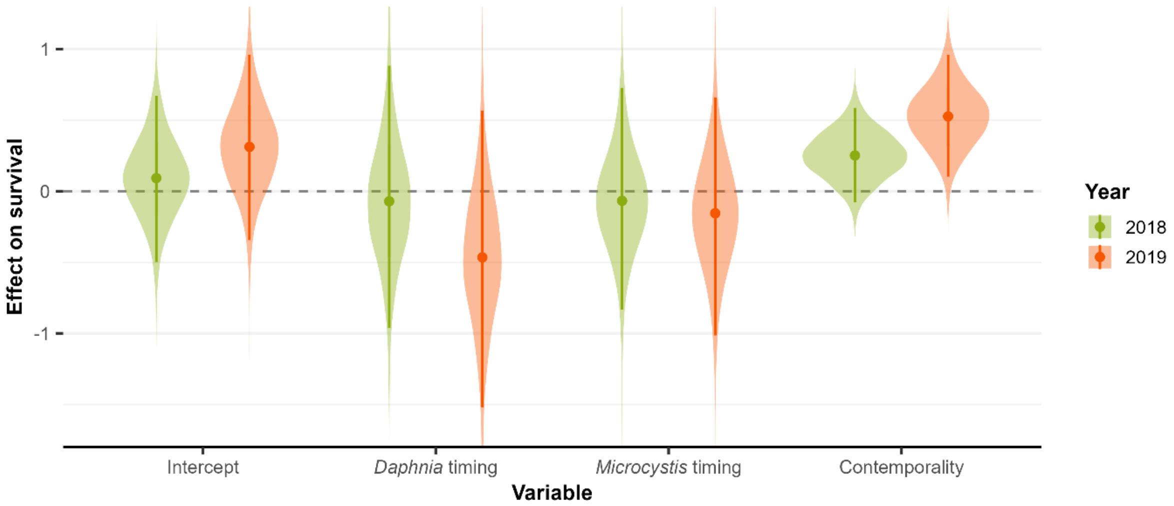


**Figure S11.** Posterior distributions of the model intercept and the effects of *Daphnia* timing, *Microcystis* timing and contemporality on *Daphnia* survival for 2018 (green) and 2019 (orange), as obtained from the interval-censored frailty survival analysis. The posterior medians and 95% credible intervals are represented by dots and vertical lines respectively.

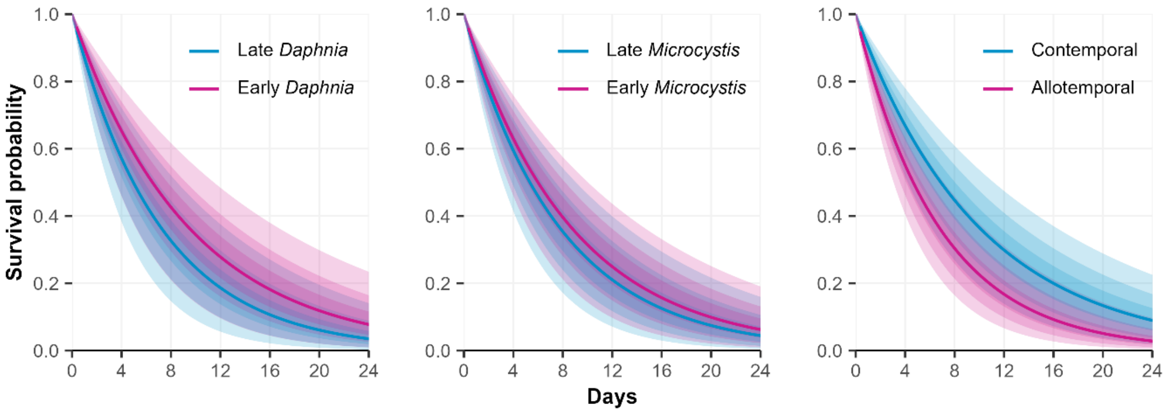


**Figure S12.** Modelled *Daphnia* survival over a period of 24 days, coloured by *Daphnia* timing (left), *Microcystis* timing (middle) and contemporality (right), as obtained from the interval-censored frailty survival analysis, assuming an exponential survival function. The median survival function is shown by a full line, while 50, 80 and 95% credible intervals are represented by means of shaded areas with an increasing transparency.

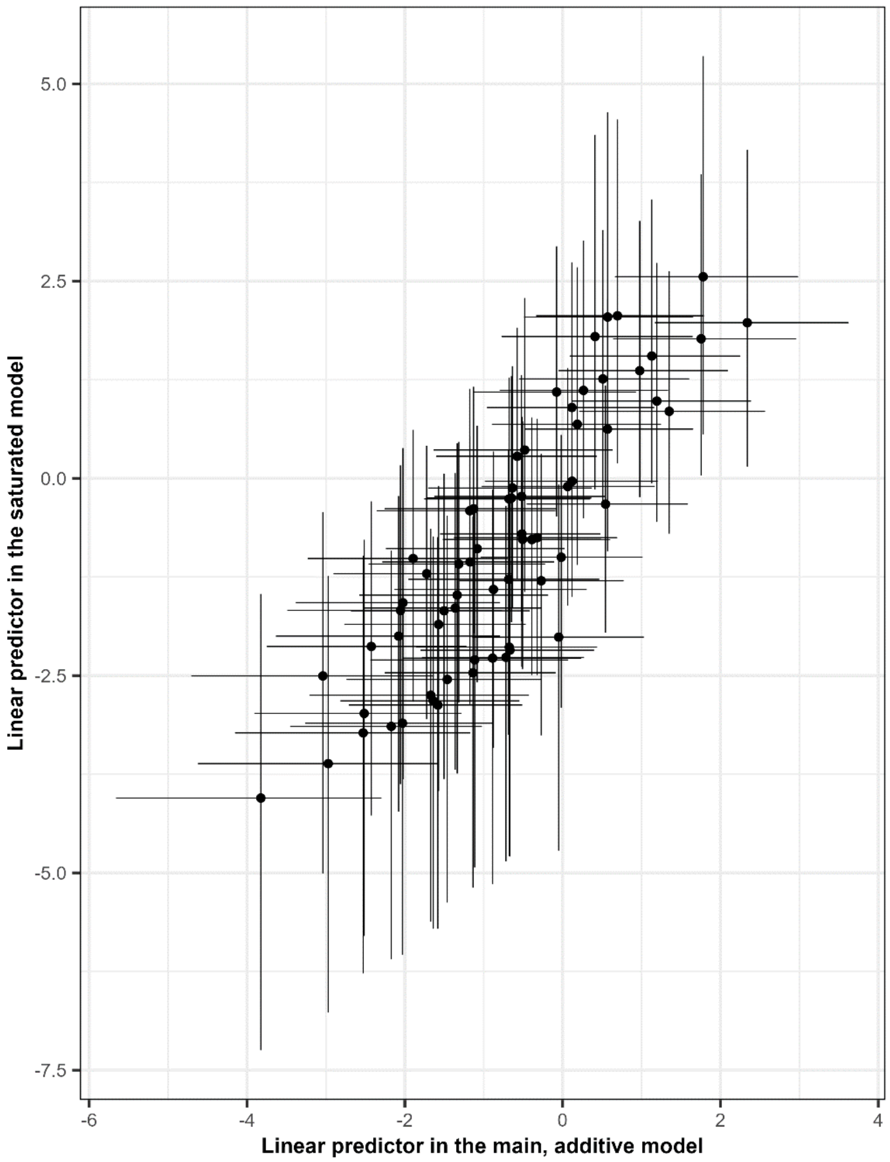


**Figure S13.** Comparison of linear predictors for the experimental units in the main model that only accounts for additive *Daphnia* and *Microcystis* genotype effects, against a fully saturated model where an additional genotype x genotype interaction effect is added. Posterior mean linear predictors and 95% credible intervals are represented by dots and horizontal or vertical lines respectively.

**Supporting tables**

**Table S1.** Sampling dates on which *D. magna* isolates and *Microcystis* sp. were isolated from Langerodevijver.

| **Timepoint** | **Sampling date** | |
|---|---|---|
| | ***D. magna*** | ***Microcystis* sp.** |
| Early 2018 | 23/04/2018 | 23/04/2018 |
| Late 2018 | 28/05/2018 | 29/05/2018 |
| Early 2019 | 25/04/2019 | 25/04/2019 |
| Late 2019 | 13/06/2019 | 26/06/2019 |

**Table S2.** Results of the year-by-year frequentist binomial GLMM model.

| Variable | Estimate | Standard error | z-value | p-value |
|---|---|---|---|---|
| **2018** | | | | |
| Intercept | -1.538 | 0.571 | -2.693 | 0.007 |
| Daphnia timing | -0.233 | 0.721 | -0.323 | 0.746 |
| Microcystis timing | 0.188 | 0.375 | 0.502 | 0.616 |
| Contemporality | 0.927 | 0.340 | 2.725 | 0.006 |
| **2019** | | | | |
| Intercept | -0.175 | 0.732 | -0.238 | 0.812 |
| Daphnia timing | -0.784 | 0.712 | -1.101 | 0.271 |
| Microcystis timing | -0.231 | 0.805 | -0.287 | 0.774 |
| Contemporality | 0.905 | 0.363 | 2.493 | 0.013 |
